# Enhanced Functional Potential of Pseudogene-associated lncRNA Genes in Mammals

**DOI:** 10.1101/2025.03.21.644463

**Authors:** Ze-Hao Zhang, Bo-Han Li, Yin-Wei Wang, Sheng Hu Qian, Lu Chen, Meng-Wei Shi, Hao Zuo, Zhen-Xia Chen

## Abstract

The functional significance of long non-coding RNAs (lncRNAs) remains a subject of debate, largely due to the complexity and cost associated with their validation experiments. However, emerging evidence suggests that pseudogenes, once viewed as genomic relics, may contribute to the origin of functional lncRNA genes. In this study spanning eight species, we systematically identified pseudogene-associated lncRNA genes using our PacBio long-read sequencing data and published RNA-seq data. Our investigation revealed that pseudogene-associated lncRNA genes exhibit heightened functional attributes compared to their non-pseudogene-associated counterparts. Notably, these pseudogene-associated lncRNAs show protein-binding proficiency, positioning them as potent regulators of gene expression. In particular, pseudogene-associated sense lncRNAs retain protein-binding capabilities inherited from parent genes of pseudogenes, thereby demonstrating greater protein-binding proficiency. Through detailed functional characterization, we elucidated the unique advantages and conserved roles of pseudogene-associated lncRNA genes, particularly in the context of gene expression regulation and DNA repair. Leveraging cross-species expression profiling, we demonstrated the prominent contribution of pseudogene-associated lncRNA genes to aging-related transcriptome changes across nine human tissues and eight mouse tissues. Overall, our findings demonstrate enhanced functional attributes of pseudogene-associated lncRNA genes and shed light on their conserved and close association with aging.

## Introduction

Long non-coding RNAs (lncRNAs), defined as RNA molecules exceeding 200 nucleotides (nt) lacking protein-coding potential, have garnered significant interest with the advances in transcriptome sequencing technology. Although tens of thousands of lncRNA genes have been identified in mammalian genomes [1], only a small fraction have been functionally characterized [2,3]. The majority of lncRNAs are still considered as transcriptional by-products [4,5], and the proportion of functional lncRNA genes in the total lncRNA gene collection remains controversial [6]. Meanwhile, function assays for lncRNA genes are usually of low throughput, high complexity and high cost, hindering comprehensive exploration of their functional roles. Pseudogenes, characterized as defective copies of protein-coding genes [7], are widely distributed in mammalian genomes [8]. Pseudogenes can be classified into multiple classes based on their mechanism of origin [9]. Duplicated pseudogenes arise from duplications at the DNA level, such as whole genome duplications and chromosomal segment duplications, whereas processed pseudogenes arise from the integration of cDNAs formed by reverse transcription of mRNAs into the genome [10]. These two classes of pseudogenes represent the two major categories of pseudogenes. Although pseudogenes exhibit sequence similarity to their corresponding parent genes, they are usually not transcribed or do not produce normal mRNA due to varying degrees of mutations in regulatory elements or coding regions [11]. Therefore, pseudogenes have long been considered as non-functional “genomic fossils” [12,13].

However, recent studies have unveiled the potential of pseudogenes as precursors for functional lncRNA genes [14]. For example, the mouse pseudogene *Rps15a-ps4* has been shown to transcribe the lncRNA Lethe [15], which regulates the transcription of downstream target genes by affecting the DNA-binding activity of the RelA subunit of the transcription factor NF-κB. The lncRNA produced by the *Oct4p4* pseudogene regulates the expression level of its parent gene *Oct4* through epigenetic modifications, thus playing an important role in stem cell differentiation [16]. In addition, lncRNAs transcribed from the *AGPG* pseudogene have been reported to promote glycolysis and cell cycle progression in human [17]. The lncRNAs produced by the pseudogenes *PCNAP1* and *PTENP1* play a role in the development of hepatocellular carcinoma and aortic aneurysm by regulating the expression levels of the parent genes *PCNA* and *PTEN*, respectively [18,19]. Together, pseudogenes can be transcribed into lncRNAs, which are involved in various regulatory functions. Accordingly, pseudogenes, which are often regarded as non-functional remnants of coding genes, may represent an appropriate substrate for the origin of functional lncRNA genes.

Here we identified pseudogene-associated lncRNA genes in vertebrates using PacBio full-length sequencing. Our findings indicate an enrichment of lncRNA genes at pseudogenes, suggesting that pseudogenes may serve as favorable substrates for the origin of functional lncRNA genes. Further characterization of pseudogene-associated lncRNA genes revealed enhanced functional attributes and protein-binding proficiency. These lncRNA genes exhibited unique advantages and conserved roles in gene expression regulation and DNA repair. Importantly, we demonstrated the prominent contribution of pseudogene-associated lncRNA genes to the aging process across nine human tissues at the single-cell level and confirmed their conserved association with aging in mouse. Taken together, our study sheds light on the heightened functional attributes of pseudogene-associated lncRNA genes and their potential implications in aging.

## Results

### Identification of pseudogene-associated lncRNA genes across eight vertebrate genomes

Long-read transcriptome sequencing has proven to be accurate and effective in improving genome annotation and discovering new genes [20,21]. To accurately characterize lncRNA genes, we performed PacBio isoform sequencing (Iso-seq), as well as RNA-seq to correct for Iso-seq data, for the adult female and male tissues (brain, heart, gut, and gonad) of three representative vertebrate models, including zebrafish (teleost), chicken (bird), and mouse (mammal) (**Figure 1**A). As expected, the transcripts detected by PacBio Iso-seq were longer in length and had a higher number of exons than the Ensembl annotated transcripts (Figure 1B, C), suggesting that our home-made datasets produced high-quality genome annotations. To broaden the scope for comparative studies, we also included five additional mammals (human, macaque, rat, rabbit, and opossum), and a total of eight species were covered for subsequent analysis. To ensure comprehensive lncRNA annotation, we integrated the Ensembl lncRNA annotation with novel lncRNAs from our home-made Iso-seq datasets and published lncRNA sets from previous studies (Methods; Figure S1), resulting in a robust annotation of 9002 (in zebrafish) to 39,461 (in human) lncRNA genes for each species (Figure 1D). Furthermore, novel lncRNA genes annotated by long-read transcriptome sequencing frequently exhibit higher transcript length, higher sequence conservation, and higher expression level in comparison with those annotated by short-read sequencing (Figure S2A–C). These results suggest that long-read transcriptome sequencing can facilitate the accurate identification of novel lncRNA genes with more functional properties.

**Figure 1.**
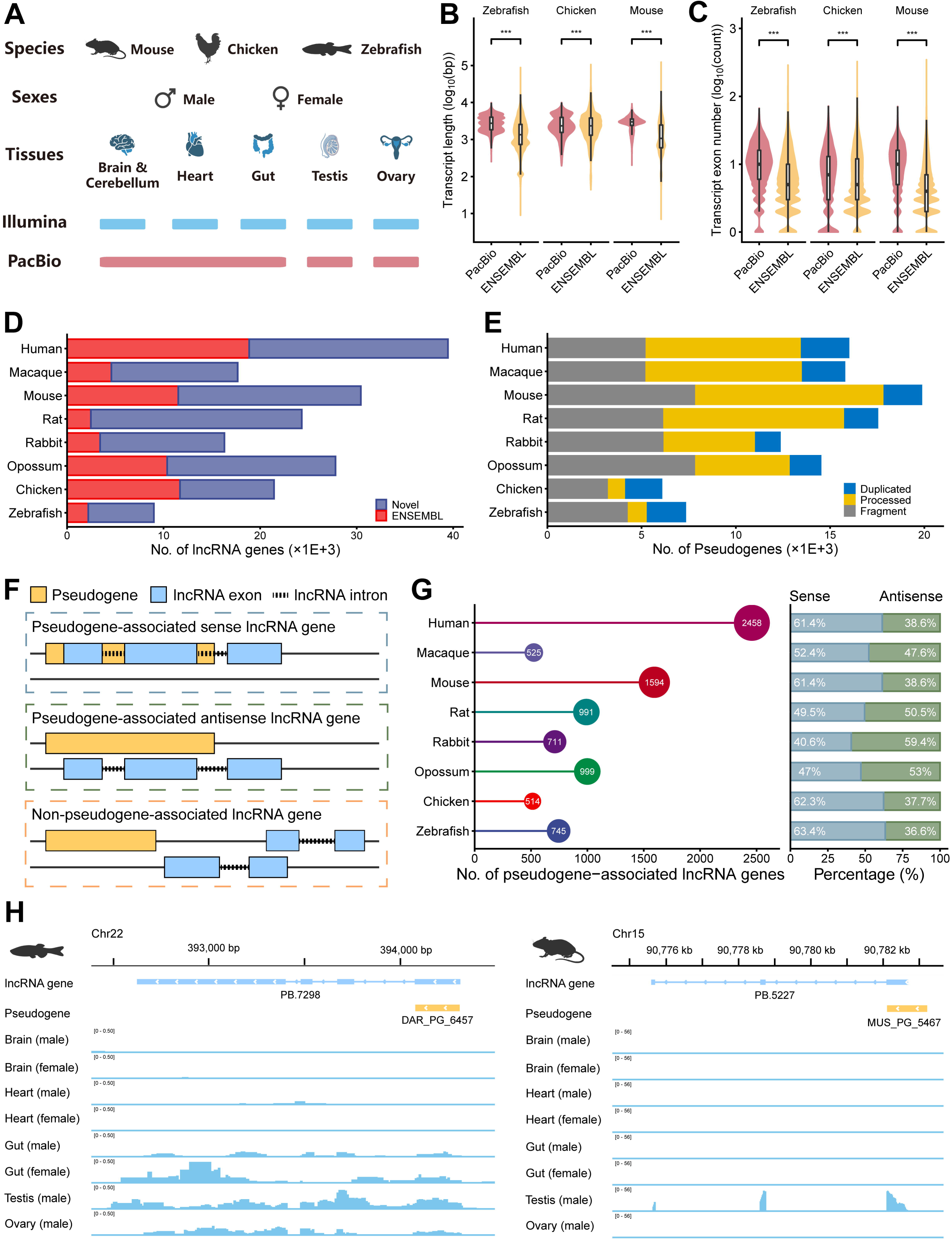
Identification of pseudogene-associated lncRNA genes. **A**. Tissue samples from zebrafish, chicken and mouse. Somatic tissues were pooled into one sample for Iso-seq library preparation for each sex. **B**. Length of transcripts in Ensembl annotation and PacBio Iso-seq. **C**. Exon number of transcripts in Ensembl annotation and PacBio Iso-seq. **D**. Number of lncRNA annotations in each species. **E**. Number of pseudogenes in each species. **F**. Methods to identify pseudogene-associated lncRNA genes. **G**. Number of pseudogene-associated lncRNA genes in each species. Bar plot showing the proportion of two categories among pseudogene-associated lncRNA genes. **H**. Expression patterns of zebrafish novel lncRNA gene *PB.7298* and mouse novel lncRNA gene *PB.5227* which are detected by Iso-seq and are identified as pseudogene-associated lncRNA genes. Iso-seq, isoform sequencing.

We then employed PseudoPipe, a computational pipeline based on homology search [22], to identify pseudogene candidates across the eight genomes. Pseudogenes that could not be reliably assessed as processed or duplicated pseudogenes were classified as fragment pseudogenes. After rigorous filtering, 6058 (in chicken) to 19,843 (in mouse) pseudogenes were retained in each species (Methods; Figure 1E). These pseudogenes covered 84.2% (in human) and 81.6% (in mouse) of the Ensembl pseudogene annotation, indicating a low false negative rate. As previous studies [8,23,24], mammals have a higher abundance of pseudogenes, with processed pseudogenes being more prevalent than duplicated pseudogenes. Conversely, in zebrafish and chicken, the proportion of duplicated pseudogenes is higher than that of processed pseudogenes (Figure S2D). Additionally, the number and density of processed pseudogenes in chicken and zebrafish are lower than in mammals (Figure S2E). Chicken CR1 reverse transcriptase is unlikely to copy polyadenylated mRNAs, whereas polyadenylated tails are potential substrates for mammalian L1 reverse transcriptase [25,26]. This may explain the differences in the abundance of processed pseudogenes between chicken and mammals. In zebrafish, the genomic coverage of retrotransposon elements is only a quarter of that in human, which could account for the lack of processed pseudogenes [27].

Next, we identified the pseudogene-associated lncRNA genes based on two different overlap patterns (Methods; Figure 1F, G; Table S1), and found 514 (in chicken) to 2458 (in human) lncRNA genes, accounting for 2.4% to 8.3% of all the lncRNA genes in each species. A robust positive correlation was identified between the number of pseudogene-associated lncRNA genes and the total number of lncRNA genes across diverse species (Figure S2F). This finding suggests that although part of the variability in the number of pseudogene-associated lncRNA genes may be biologically meaningful, much is likely to be explained by unequal size of the lncRNA gene repertoire. Consequently, factors influencing the size of the lncRNA annotation, including the quality of genome sequence and annotation, as well as the amount and sequencing depth of RNA-seq data used to identify novel lncRNA genes [28], affect the number of pseudogene-associated lncRNA genes. Additionally, 41.7% (in human) to 75.2% (in rat) of the pseudogene-associated lncRNA genes in each species were novel lncRNA genes (Figure S2G). This highlights the significance of comprehensive lncRNA annotation. For example, zebrafish novel lncRNA gene *PB.7298* and mouse novel lncRNA gene *PB.5227* detected by Iso-seq overlapped with the pseudogene and were identified as pseudogene-associated lncRNA genes, with the first exon of both derived from the pseudogene (Figure 1H). *PB.7298* was found to be expressed in the gut and gonads, whereas *PB.5227* was only expressed in the testis.

### Enrichment of lncRNA genes at pseudogenes across species

To understand the contribution of pseudogenes to the makeup and function of lncRNA genes, we evaluated the enrichment of lncRNA genes at pseudogenes (Methods) and found that both sense and antisense lncRNA genes were enriched at pseudogenes compared to shuffled regions in mammals and zebrafish (empirical *P* ≤ 0.001) (**Figure 2**A). After excluding the potential effects of chromosomal biases or repetitive sequences (Methods), similar results were still observed (Figure 2A). We further evaluated the enrichment of lncRNA genes at each class of pseudogenes, and found that both duplicated and processed pseudogenes showed high enrichment of lncRNA genes in comparison to shuffled regions (Figure S3). These findings imply that pseudogenes act as relics of protein-coding gene copies to provide favorable substrates for the origin of functional lncRNA genes, which are retained by purifying selection during evolution. To further confirm whether the origin of lncRNA genes is functionally gained or lost for pseudogene regions. We evaluated the sequence conservation of overlapping and non-overlapping regions within the pseudogenes overlapping with sense and antisense lncRNA genes using PhastCons score in human and mouse. We found that regions within pseudogenes that overlap with exons of lncRNA genes are more conserved than non-overlapping portions (two-sided Wilcoxon rank sum test, \**P* < 0.05, \*\*\**P* < 0.001) (Figure 2B), suggesting that these pseudogene regions acquire enhanced functional properties through the origin of sense or antisense lncRNA genes. For example, within the pseudogene HOS_PG_6183, which overlaps with the human lncRNA gene *ROCK1P1*, both PhastCons and PhyloP scores showed higher sequence conservation in the regions overlapping the *ROCK1P1* exons (Figure 2C).

**Figure 2.**
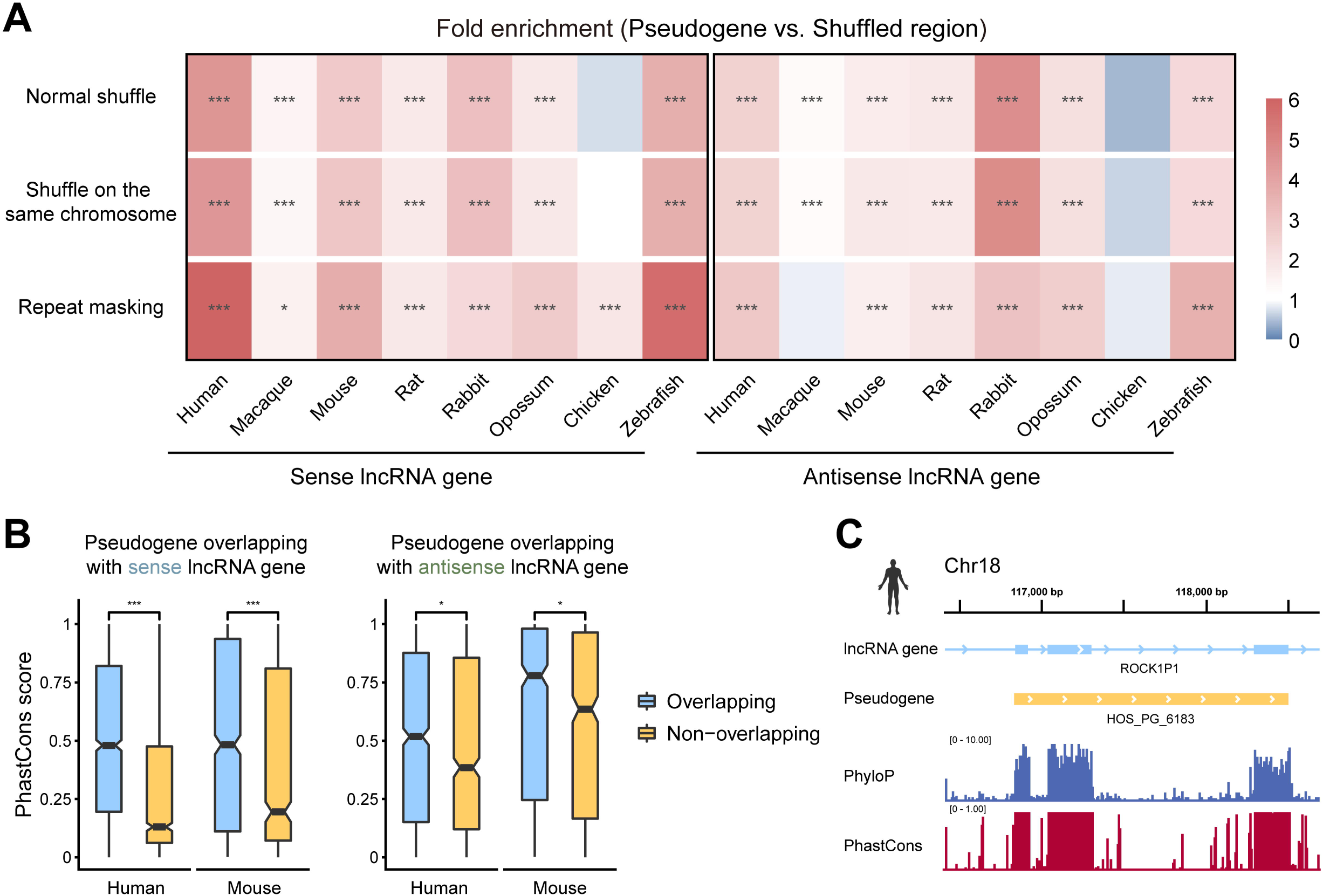
lncRNA genes are enriched at pseudogenes. **A**. Enrichment of sense and antisense lncRNA genes at pseudogenes in each species. \**P* ≤ 0.05, \*\**P* ≤ 0.01, \*\*\**P* ≤ 0.001. **B**. Sequence conservation (PhastCons score), for overlapping and non-overlapping regions within the pseudogenes overlapping with sense and antisense lncRNA genes in human and mouse. **C**. Genomic profile depicts a high conservation of regions within the pseudogene HOS_PG_6183 overlapping the *ROCK1P1* exons. The per-base phastCons, and the phyloP conservation scores are presented.

### Enhanced functional attributes of pseudogene-associated lncRNA genes in sequence

The enrichment of lncRNA genes at pseudogenes implies that pseudogenes may have endowed pseudogene-associated lncRNA genes with functionality. To explore the functional potential of pseudogene-associated lncRNA genes, we conducted comparative analyses with non-pseudogene-associated (NPA) counterparts, focusing on evolutionary age, sequence conservation, intrapopulation variation, and structural attributes.

We estimated the evolutionary age of human and mouse lncRNA genes using 1:1 orthologous lncRNA genes based on the phylogenetic relationships of each species (Methods; Table S2). Although large proportions of lncRNA genes were species-specific in human (38.9%) and mouse (35.6%), both pseudogene-associated sense (PAS) and pseudogene-associated antisense (PAA) lncRNA genes had higher proportions of genes in older age groups, and lower proportions of genes in younger age groups than NPA lncRNA genes (**Figure 3**A; Figure S4A). Next, we analyzed the enrichment of age groups for each category of lncRNA genes using all lncRNA genes as background genes. Take human for example, we found that NPA lncRNA genes were not enriched in any of the evolutionary age groups (Figure 3B). In contrast, PAS lncRNA genes were enriched in 429 million years ago (Ma), and PAA lncRNA genes were enriched in 429 Ma, 319 Ma, and 160 Ma. A similar phenomenon was observed in mouse (Figure S4B). Taken together, our analysis suggests that pseudogene-associated lncRNA genes exhibit higher evolutionary conservation, which implies their greater functionality.

**Figure 3.**
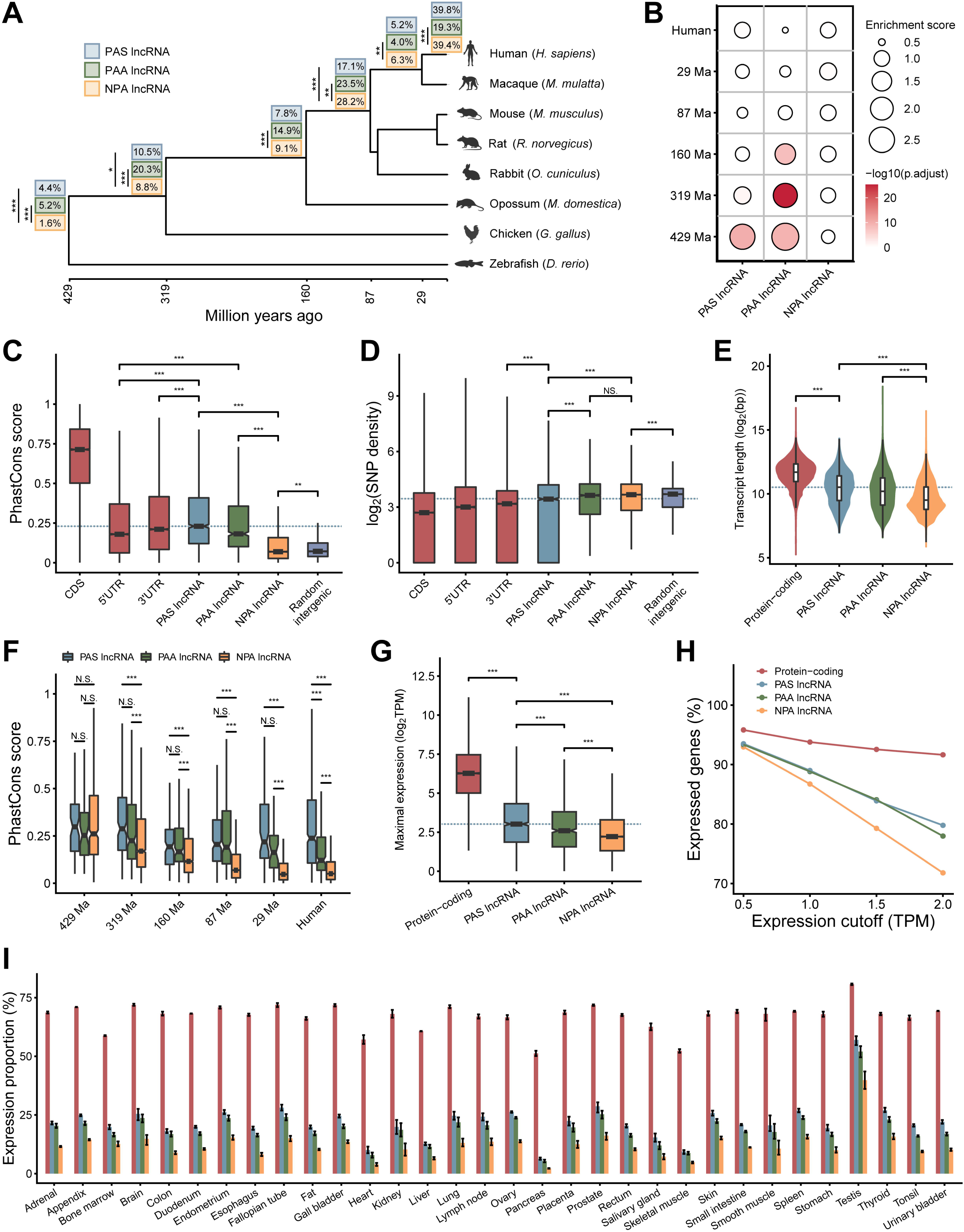
Functionality characterization and expression profiles of pseudogene-associated lncRNA genes. **A**. Distribution of PAS lncRNA genes, PAA lncRNA genes, and NPA lncRNA genes in the branches of the human phylogenetic tree. Tree tips, lncRNA proportions for each age group. \**P* < 0.05, \*\**P* < 0.01, \*\*\**P* < 0.001, two-sided Fisher’s exact test. **B**. Enrichment of PAS lncRNA genes, PAA lncRNA genes, and NPA lncRNA genes at different evolutionary ages in human. A total of 2000 randomly selected NPA lncRNA genes were utilized as controls. **C**. Exonic sequence conservation (PhastCons score), for CDS, UTR, PAS lncRNA genes, PAA lncRNA genes, NPA lncRNA genes, and random intergenic regions in human. **D**. Exon common SNP density of CDS, UTR, PAS lncRNA genes, PAA lncRNA genes, NPA lncRNA genes, and random intergenic regions in human. **E**. Transcript lengths of protein-coding genes, PAS lncRNA genes, PAA lncRNA genes, and NPA lncRNA genes in human. **F**. Exonic sequence conservation (PhastCons score), for PAS lncRNA genes, PAA lncRNA genes, and NPA lncRNA genes in each age group in human. **G**. Maximum expression levels of protein-coding genes, PAS lncRNA genes, PAA lncRNA genes, and NPA lncRNA genes in human. **H**. Expression proportions of protein-coding genes, PAS lncRNA genes, PAA lncRNA genes, and NPA lncRNA genes at a series of expression thresholds in human. **I**. Mean expression proportions of protein-coding genes, PAS lncRNA genes, PAA lncRNA genes, and NPA lncRNA genes in various tissues in human. Error bars indicate standard deviation of the proportion of expression. Legend is the same as in Figure 3H. PAS, pseudogene-associated sense; PAA, pseudogene-associated antisense; NPA, non-pseudogene-associated; Ma, million years ago; CDS, coding sequence; UTR, untranslated region; SNP, single nucleotide polymorphism.

To further reveal the functional potential of pseudogene-associated lncRNA genes, we evaluated the cross-species DNA sequence conservation of these lncRNA genes (Figure 3C; Figure S4C; Table S2). We found that the exon sequence conservation of both PAS (median ∼0.23 in human and ∼0.24 in mouse) and PAA lncRNA genes (median ∼0.18 in human and ∼0.29 in mouse) was lower than that of the coding sequence (CDS) of protein-coding genes, but higher than that of NPA lncRNA genes (median ∼0.07 in human and ∼0.11 in mouse) and random intergenic regions (median ∼0.07 in human and ∼0.09 in mouse), and even higher than that of the 5’ untranslated regions (UTRs) of protein-coding genes in both human and mouse (two-sided Wilcoxon rank sum test, *P* < 0.001). Conservation beyond the non-coding regions of the exons of protein-coding genes, demonstrating greater functionality of pseudogene-associated lncRNA genes. To ensure that the higher conservation of pseudogene-associated lncRNA genes compared to NPA counterparts was not due to their older age, we evaluated the conservation of lncRNA genes in each age group (Figure 3F; Figure S4D). We found that each category of lncRNA genes is highly conserved in the oldest age group, but both PAS and PAA lncRNA genes remain consistently more conserved than NPA counterparts in other age groups, suggesting stronger evolutionary constraints on pseudogene-associated lncRNA genes at each age.

Next, we investigated the intrapopulation variation of pseudogene-associated lncRNA genes in human [29]. Although the derived allele frequencies of single nucleotide polymorphisms (SNP) found in the exons of pseudogene-associated lncRNA genes were not different from those found in the exons of NPA lncRNA genes (two-sided unpaired T test, *P* > 0.05) (Figure S4E), the SNP density of exons of PAS lncRNA genes was lower than that of exons of NPA lncRNA genes as well as random intergenic regions (two-sided Wilcoxon rank sum test, *P* < 0.001) (Figure 3D). This suggests that PAS lncRNA genes may be subject to slightly stronger intrapopulation purifying selection than NPA lncRNA genes. In addition, the Combined Annotation-Dependent Depletion (CADD) score [30] is a deleteriousness score for human variants. We found that CADD scores for SNPs in exons of both PAS and PAA lncRNA genes were higher than those in exons of NPA lncRNA genes (two-sided Wilcoxon rank sum test, *P* < 0.001) (Figure S4F), suggesting that variants in pseudogene-associated lncRNA genes are more pathogenic than those in NPA lncRNA genes. These results again implied the potential functionality of pseudogene-associated lncRNA genes.

Since the functionality of genes is often related to the structural attributes of their transcripts [1,31], we examined the structural attributes of the transcripts of pseudogene-associated lncRNA genes (Table S2). We found that both PAS and PAA lncRNA genes had higher transcript lengths and exon numbers than NPA lncRNA genes (two-sided Wilcoxon rank sum test, *P* < 0.001; two-sided unpaired T test, *P* < 0.001) (Figure 3E; Figure S4G, H), suggesting that the transcripts of pseudogene-associated lncRNA genes may carry more RNA structural domains, thereby favoring their interactions with other nucleic acid or protein molecules [32].

Altogether, these findings in sequence revealed that pseudogene-associated lncRNA genes possess enhanced functional attributes compared to NPA counterparts.

### Heightened functional attributes of pseudogene-associated lncRNA genes in expression

To further explore the functionality of pseudogene-associated lncRNA genes in expression, we conducted comparisons of their expression profiles and epigenetic features, including chromatin accessibility, histone modifications, and DNA methylation, near transcription start sites (TSSs) with protein-coding genes and NPA lncRNA genes.

Utilizing RNA-seq data from 32 human tissues, we determined expression profiles across tissues and found that the expression level of both PAS and PAA lncRNA genes, compared with NPA lncRNA genes, was closer to protein-coding genes (two-sided Wilcoxon rank sum test, *P* < 0.001) (Figure 3G; Figure S5; Table S2). Moreover, the expression ratios of both PAS and PAA lncRNA genes were higher than that of NPA lncRNA genes, and closer to that of protein-coding genes at all expression cutoffs (Figure 3H) and in all 32 tissues (Figure 3I). Notably, PAS lncRNA genes show the highest expression activity. We also observed a conserved high expression activity of PAS lncRNA genes in mouse (Figure S6A-D). In addition, each category of lncRNA genes both showed tissue-specific expression patterns (Figure S6E; Table S2), supporting the high tissue-specificity of lncRNA gene expression [33,34]. Meanwhile, both PAS and PAA lncRNA genes (median: 7 and 6 tissues), as well as coding genes, exhibited broader tissue expression profiles than NPA lncRNA genes (median: 4 tissues) in human (two-sided Wilcoxon rank sum test, *P* < 0.001) (Figure S6E), demonstrating their potential functional versatility across tissues.

Chromatin accessibility plays a crucial role in regulating gene expression by influencing the accessibility of DNA to transcriptional machinery and regulatory proteins. Therefore, to elucidate the causes of active expression of pseudogene-associated lncRNA genes at the epigenomic level, we characterized chromatin accessibility near the transcription start sites (TSSs) in human. Compared to NPA lncRNA genes, both PAS and PAA lncRNA genes showed more housekeeping chromatin open regions by ATAC-seq, and PAS lncRNA genes also showed higher DNase accessibility by DNase-seq (**Figure 4**A, B), demonstating higher chromatin accessibility at proximal promoters of pseudogene-associated lncRNA genes.

**Figure 4.**
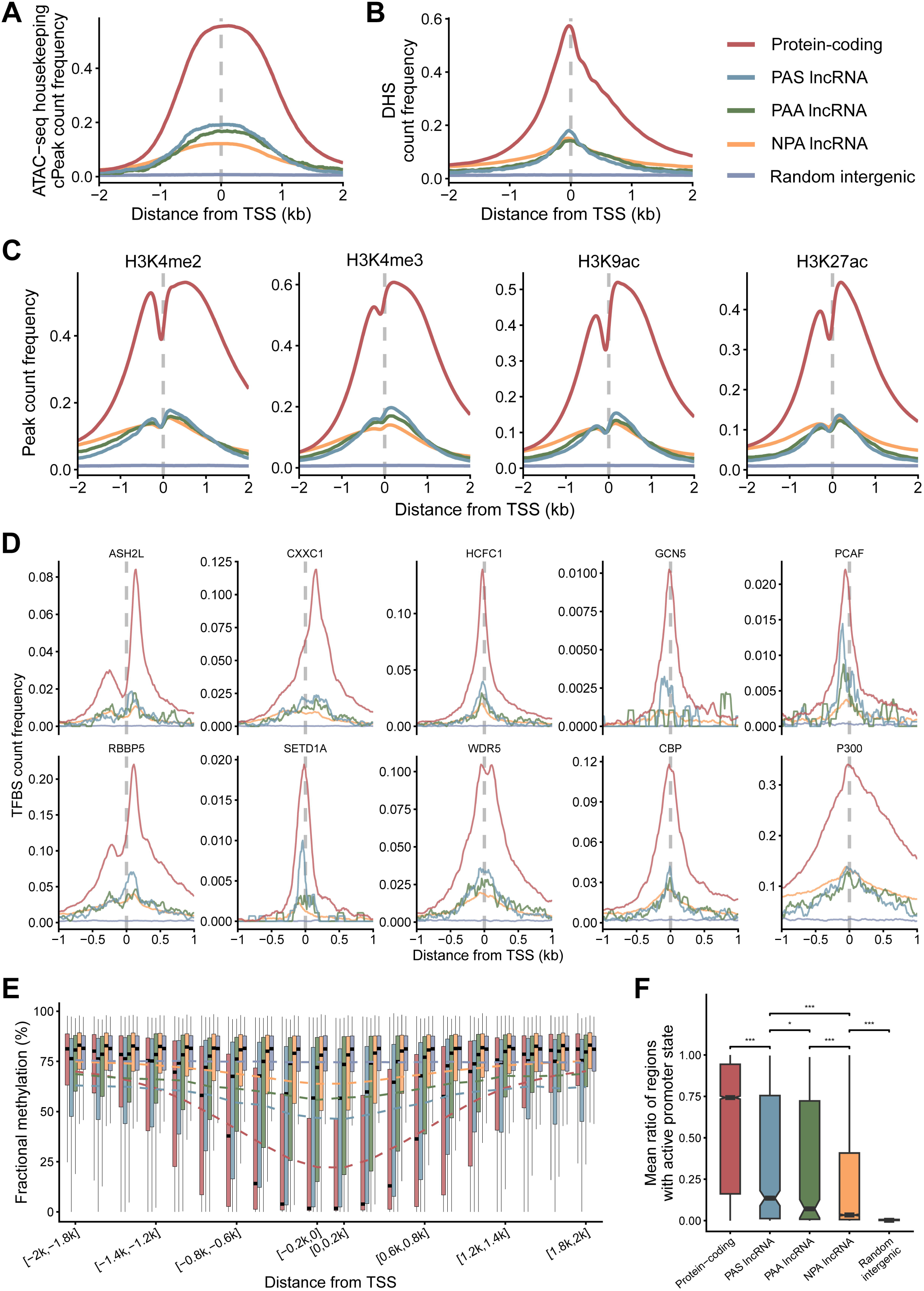
Epigenomic features of pseudogene-associated lncRNA genes. **A**. Distribution of housekeeping consensus peaks from ATAC-seq in a total of 4kb regions upstream and downstream of protein-coding gene, PAS lncRNA genes, PAA lncRNA genes, and NPA lncRNA genes TSSs, and random intergenic regions. **B**. Distribution of DHS in a total of 4kb regions upstream and downstream of protein-coding gene, PAS lncRNA genes, PAA lncRNA genes, and NPA lncRNA genes TSSs, and random intergenic regions. **C**. Distribution of H3K4me3, H3K4me2, H3K9ac, and H3K27ac peaks in a total of 4kb regions upstream and downstream of protein-coding gene, PAS lncRNA genes, PAA lncRNA genes, and NPA lncRNA genes TSSs, and random intergenic regions. **D**. Distribution of ten chromatin modifiers binding sites in a total of 2kb regions upstream and downstream of protein-coding gene, PAS lncRNA genes, PAA lncRNA genes, and NPA lncRNA genes TSSs, and random intergenic regions. **E**. DNA methylation in each 200bp window within a total of 4kb regions upstream and downstream of protein-coding gene, PAS lncRNA genes, PAA lncRNA genes, and NPA lncRNA genes TSSs, and random intergenic regions. Dotted lines indicate average DNA methylation values over the entire region. Legend is the same as in Figure 4A. **F**. Ratios of active promoter chromHMM state regions in the proximal promoters of protein-coding genes, PAS lncRNA genes, PAA lncRNA genes, NPA lncRNA genes, and random intergenic regions. TSS, transcription start site; DHS, DNase I hypersensitivity site.

Histone modifications may regulate gene expression by influencing chromatin structure and function. We found more signals near TSS of both PAS and PAA lncRNA genes by ChIP-seq for histone modifications associated with open chromatin, including H3K4me3, H3K4me2, and H3K9ac, than that near TSS of NPA lncRNA genes (Figure 4C). PAS lncRNA genes also exhibited slightly higher H3K27ac signals compared to NPA lncRNA genes. These results are consistent with the higher chromatin accessibility near pseudogene-associated lncRNA genes. Moreover, we counted around the TSS the binding sites of ten chromatin modifying factors, including six components of the H3K4 methyltransferase Complex of Proteins Associated with Set1 (COMPASS) [35], two components of the histone acetyltransferase CBP/p300 complex [36], and two componets of the histone acetyltransferase GCN5/PCAF complex [37], and found that the binding levels of all the chromatin modifying factors except p300 were elevated in the proximal regions of both PAS and PAA lncRNA gene TSSs (Figure 4D), indicating the shaping of epigenomic signatures characteristic of enhanced transcriptional regulation. The higher signials of histone modifications associated with open chromatin and the higher binding levels to related chromatin modifiers near the TSS of pseudogene-associated lncRNA genes would allow their transcriptional activation.

Besides histone modifications, DNA methylation are also important regulators of gene expression. We thus examined DNA methylation in each 200 bp window within a total of 4kb regions upstream and downstream of the gene TSSs, and found that the DNA methylation of both PAS and PAA lncRNA genes was lower than that of NPA lncRNA genes in each window (two-sided Wilcoxon rank sum test, *P* < 0.001) (Figure 4E), and reached the lowest level at TSS, indicating that the pseudogene-associated lncRNA genes could more effectively avoid the silencing of gene expression caused by DNA methylation in the promoter region.

As an integration of chromatin accessibility, histone modifications, DNA methylation, and other epigenetic feature, chromHMM chromatin states serve as a molecular framework influencing gene expression patterns [38]. We found that the ratio of active promoter state regions in the proximal promoters of both PAS and PAA lncRNA genes was higher than that in the proximal promoters of NPA lncRNA genes as well as in the random intergenic region (two-sided Wilcoxon rank sum test, *P* < 0.001) (Figure 4F). These results are consistent with their epigenomic characterization.

In summary, the characterization of gene expression, chromatin accessibility, histone modifications, DNA methylation levels, and chromatin state collectively indicated that pseudogene-associated lncRNA genes are subject to heightened levels of active transcriptional regulation, thereby facilitating enhanced functionality.

### Protein-binding proficiency of pseudogene-associated lncRNAs hints they are important regulators of gene expression

lncRNAs usually function as regulators of gene expression [39]. In general, lncRNAs often exert their regulatory roles by interacting with RNA-binding proteins (RBPs) [40,41]. They regulate gene expression at the transcriptional level by recruiting transcription factors and chromatin modifiers, or at the post-transcriptional level by competing with mRNAs for binding to RNA stabilizing/destabilizing factors [2,42,43]. Therefore, to explore the potential function of pseudogene-associated lncRNA genes, we scrutinized their features associated with RNA-protein binding in human.

N6-methyladenosine (m6A) is a prevalent and significant RNA modification known to influence the binding between RNAs and proteins [44,45]. We investigated the m6A modification in pseudogene-associated lncRNAs and found that the density of m6A sites in both PAS and PAA lncRNAs was higher than that in NPA lncRNAs (two-sided Wilcoxon rank sum test, *P* < 0.001) (**Figure 5**A; Table S2), suggesting a more complex binding pattern between pseudogene-associated lncRNAs and proteins. Next, we directly characterized the binding of pseudogene-associated lncRNAs to RBPs (Table S2). In transcripts with RBP binding sites, the richness of RBPs bound to both PAS and PAA lncRNAs was higher than those bound to NPA lncRNAs (two-sided Wilcoxon rank sum test, \*\**P* < 0.01, \*\*\**P* < 0.001) (Figure 5B). Furthermore, the diversity of RBPs bound to PAS lncRNAs was also higher than those bound to NPA lncRNAs (Figure S7A). To exclude the effect of gene expression levels on the detection of m6A sites as well as RBP binding sites, we evaluated these features associated with RNA-protein binding in four expression level groups and found similar results (Figure S7B-D). These results suggest that pseudogene-associated lncRNAs bind more abundant RBPs than NPA lncRNAs, indicating that pseudogene-associated lncRNAs are greater regulators of gene expression compared to NPA counterparts.

**Figure 5.**
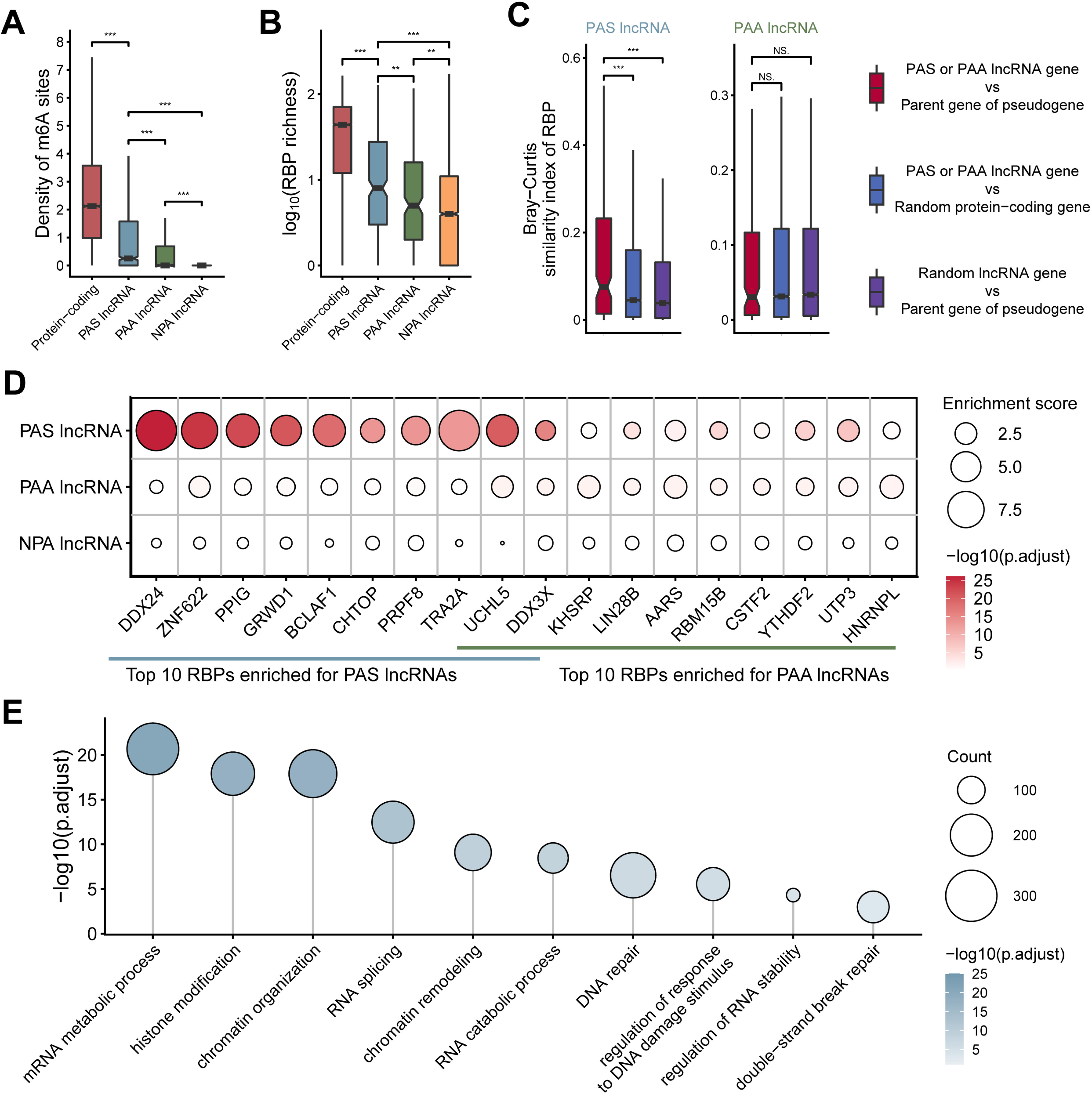
RNA-protein binding related features and putative functions of pseudogene-associated lncRNA genes. **A**. Density of m6A sites in protein-coding gene transcripts, PAS lncRNAs, PAA lncRNAs, and NPA lncRNAs. **B**. Richness of RBPs bound to protein-coding gene transcripts, PAS lncRNAs, PAA lncRNAs, and NPA lncRNAs. **C**. Bray-Curtis similarity between RBPs bound to pseudogene-associated lncRNAs and RBPs bound to transcripts of parent genes of pseudogenes. Two types of gene pairs were utilized as controls: pseudogene-associated lncRNA genes and random protein-coding genes that exhibited a matched expression level with parent genes of pseudogenes, as well as parent genes of pseudogenes and random lncRNA genes that exhibited a matched expression level with pseudogene-associated lncRNA genes. **D**. Top 10 RBPs enriched for PAS and PAA lncRNAs. A total of 2000 randomly selected NPA lncRNAs were used as controls. **E**. GO enrichment analysis to annotate the function of PAS lncRNA genes in human. The representative GO biological processes are shown. m6A, N6-methyladenosine; RBP, RNA-binding protein; GO, Gene Ontology.

Although both PAS and PAA lncRNAs showed advantages in protein-binding capacity, PAS lncRNAs exhibited higher m6A site density as well as more abundant and diverse RBP binding sites compared to PAA lncRNAs (Figure 5A, B; Figure S7A). Since PAS lncRNAs have homologous sequences of transcripts from parent genes of pseudogenes, whereas PAA lncRNAs have potential complementary sequences of transcripts from parent genes of pseudogenes, we hypothesized that the abundant RBP binding sites of PAS lncRNAs are conferred by parent genes of pseudogenes. To test this hypothesis, we calculated the Bray-Curtis similarity of RBP binding sites between each category of pseudogene-associated lncRNAs and transcripts from parent genes of pseudogenes. A higher RBP similarity was identified between PAS lncRNAs and transcripts from parent genes of pseudogenes compared to gene pairs used as controls (two-sided Wilcoxon rank sum test, *P* < 0.001) (Figure 5C). However, the RBP similarity between PAA lncRNAs and transcripts from parent genes of pseudogenes did not differ from that observed in gene pairs used as controls (Figure 5C). These findings suggest that only PAS lncRNAs inherit the RBP binding sites from parent genes of pseudogenes, thereby demonstrating greater proficiency in protein-binding.

Furthermore, we analyzed the enrichment of RBPs for each category of lncRNAs using all lncRNAs as background. We found that NPA lncRNAs were not enriched for RBPs, whereas PAS lncRNAs and PAA lncRNAs were enriched for 104 and 12 RBPs, respectively (Figure S7E). These RBPs include a number of gene expression regulators (Figure 5D). The top 10 RBPs enriched by PAS lncRNAs include transcriptional regulators (ZNF662, GRWD1, BCLAF1, CHTOP, DDX3X), RNA splicing regulators (PPIG, PRPF8, TRA2A), deubiquitinase (UCHL5), and mRNA stabilizing factor (DDX24). Meanwhile, PAA lncRNAs were also enriched for DDX3X and UCHL5, as well as other gene expression regulators such as regulators of RNA splicing and processing (RBM15B, CSTF2, HNRNPL), translation regulator (LIN28B), and mRNA degradation factors (KHSRP, YTHDF2). These results suggest that pseudogene-associated lncRNAs may be involved in the regulation of gene expression at multiple levels by binding to RBPs. Moreover, DDX24 [46] and DDX3X [47,48] were found to be closely associated with DNA repair, indicating a potential involvement of pseudogene-associated lncRNAs in DNA repair.

Co-expression can indicate functional relevance or regulatory relationships between lncRNAs and protein-coding genes [28]. To infer the putative functions of pseudogene-associated lncRNA genes, we identified co-expressed protein-coding genes for each lncRNA gene in human and mouse (Methods). We further analyzed the enrichment of co-expressed genes to identify co-expressed genes more closely associated with the function of each category of lncRNA genes. We found that both PAS and PAA lncRNA genes were enriched for more co-expressed protein-coding genes compared to NPA lncRNA genes in both human and mouse (Figure S7F, G). Next, co-expressed genes enriched by each category of lncRNA genes were analyzed for Gene Ontology (GO) enrichment analysis. The co-expressed genes enriched by NPA lncRNA genes were not enriched in any of the entries in human and were enriched in mitosis-related entries in mouse (Figure S7H). Whereas the co-expressed genes enriched by PAS lncRNA genes were enriched in functions related to the regulation of gene expression, such as histone modification, chromatin organization, chromatin remodeling, mRNA metabolic process, RNA catabolic process, and regulation of RNA stability in both human and mouse (Figure 5E; Figure S7I). Meanwhile, the co-expressed genes enriched by PAA lncRNA genes were enriched in similar functions related to gene expression regulation in mouse (Figure S7J, K). These results again implies the important role of pseudogene-associated lncRNA genes in gene expression regulation. Enrichment analysis also revealed that pseudogene-associated lncRNA genes have conserved functions in DNA repair, double-strand break repair, and regulation of response to DNA damage stimulus (Figure 5E; Figure S7I, K).

Taken together, our analyses about RBPs and co-expression collectively suggested that pseudogene-associated lncRNA genes have unique advantages and conserved functions in gene expression regulation and DNA repair, which may explain the high conservation of pseudogene-associated lncRNA genes.

### Deciphering aging-related expression changes of pseudogene-associated lncRNA genes across tissues

Aging is often accompanied by reprogramming of the transcriptome and downregulation of DNA repair levels. Motivated by the putative functions of pseudogene-associated lncRNA genes, we extended our scope to aging. To comprehensively explore the aging-related expression alterations of pseudogene-associated lncRNA genes, we used single-cell/nucleus RNA sequencing (sc/snRNA-seq) data from 9 human tissues to characterize the expression profile of pseudogene-associated lncRNA genes in young and old groups (Methods; Table S3).

After cell filtration, we retained a total of 850,860 high-quality single cells/nuclei and identified 72 cell types based on canonical cell markers in all tissues (Figure S8; Table S4). We found that both PAS and PAA lncRNA genes were more actively expressed than NPA lncRNA genes in each cell type of each tissue (two-sided Fisher’s exact test, *P* < 0.001) (Figure S9, 10). Meanwhile, both PAS and PAA lncRNA genes, along with coding genes, displayed more extensive cell type expression profiles than NPA lncRNA genes across all tissues (two-sided Wilcoxon rank sum test, *P* < 0.001) (Figure S11), indicating their potential functional versatility across cell types.

Subsequently, a differential expression analysis was conducted between the young and old age groups in each cell type of each tissue (Table S5). We found that the proportion of aging-related differentially expressed genes (DEGs) in both PAS and PAA lncRNA genes was higher than that in NPA lncRNA genes in each tissue (**Figure 6**A, B), suggesting a more prominent contribution of pseudogene-associated lncRNA genes to aging-related transcriptome changes and their closer association with aging. Furthermore, we also observed that the close association of pseudogene-associated lncRNA genes with aging was cell type-specific (Figure S12). For instance, in the brain, the proportion of up-regulated genes in PAS lncRNA genes was higher than that in NPA lncRNA genes in 6 out of 8 cell types, suggesting that pseudogene-associated lncRNA genes respond differently to aging in different cell types of the same tissue. This underscores the necessity for a detailed examination of their aging-related alterations in expression at the single-cell level.

**Figure 6.**
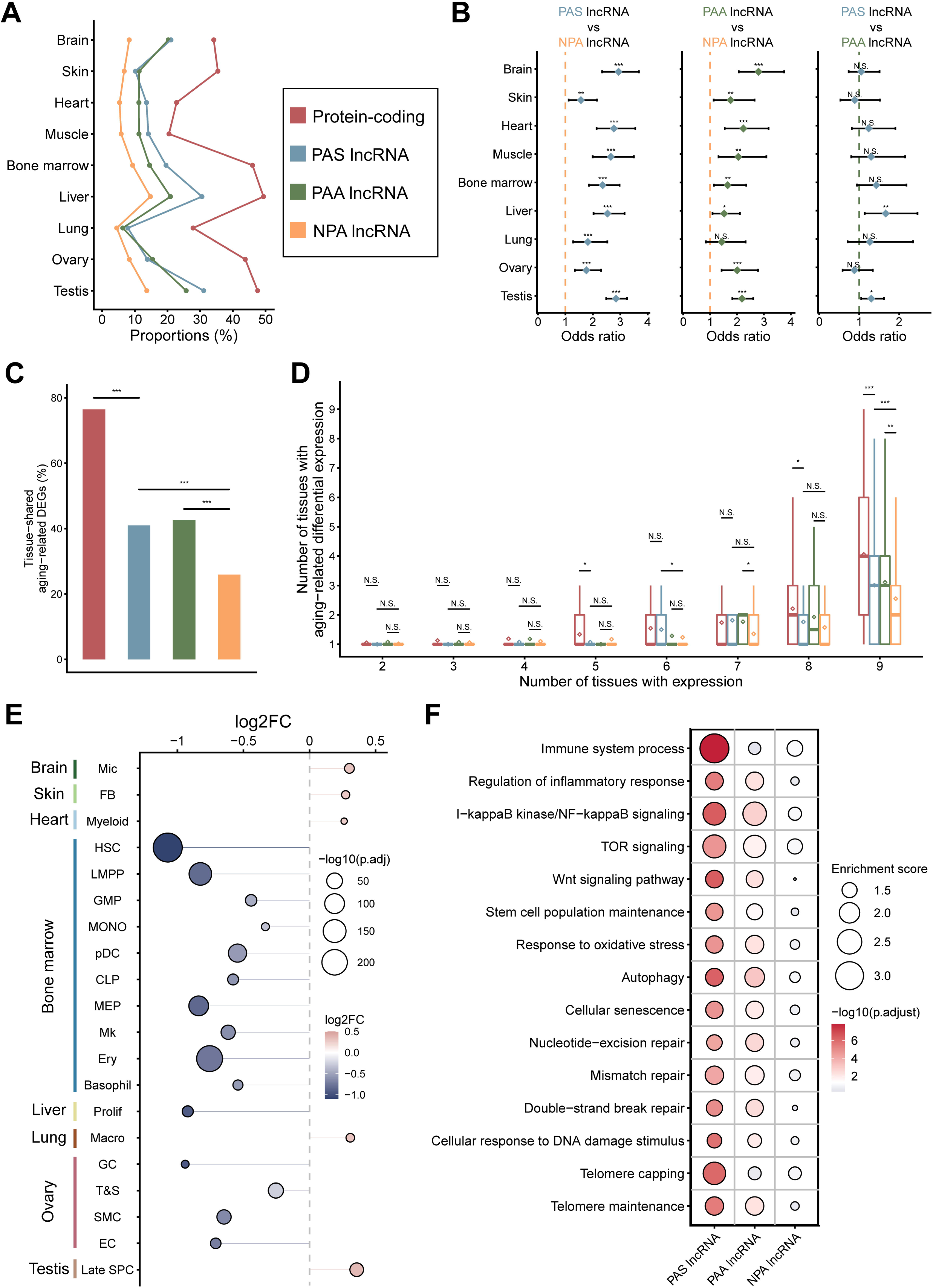
Aging-related expression changes of pseudogene-associated lncRNA genes across nine human tissues. **A**. Proportions of aging-related DEGs for protein-coding genes, PAS lncRNA genes, PAA lncRNA genes, and NPA lncRNA genes in each tissue. **B**. Odds ratios and confidence intervals (95%) between the proportions of aging-related DEGs in each category of lncRNA genes in nine tissues. \**P* < 0.05, \*\**P* < 0.01, \*\*\**P* < 0.001, two-sided Fisher’s exact tests were performed for comparisons between each category of lncRNA genes. **C**. Among aging-related DEGs, proportions of tissue-shared aging-related DEGs for protein-coding genes, PAS lncRNA genes, PAA lncRNA genes, and NPA lncRNA genes. **D**. Number of tissues in which the genes exhibited aging-related differential expression for protein-coding genes, PAS lncRNA genes, PAA lncRNA genes, and NPA lncRNA genes in each expression breadth group. The diamond dots represent the average. \**P* < 0.05, \*\**P* < 0.01, \*\*\**P* < 0.001, two-sided Wilcoxon rank sum test. **E**. Aging-related differential expression of *JPX* in each cell type of each tissue. **F**. GO enrichment analysis based on guilt-by-association method to annotate the function of aging-related DEGs in each category of lncRNA genes. The representative GO biological processes are shown. A total of 2000 randomly selected NPA lncRNA genes were used as controls for comparison. DEG, differentially expressed gene; Mic, microglia; FB, fibroblast; Myeloid, myeloid cell; HSC, hematopoietic stem cell; LMPP, lympho-myeloid primed progenitor; GMP, granulocyte-monocyte progenitor; MONO, monocyte; pDC, plasmacytoid dendritic cell; CLP, common lymphoid progenitor; MEP, megakaryocyte-erythroid progenitor; Mk, megakaryocyte; Ery, erythroid; Basophil, basophil cell; Prolif, proliferative cell; Macro, macrophage; GC, granulosa cell; T&S, theca and stroma cell; SMC, smooth muscle cell; EC, endothelial cell; Late SPC, late primary spermatocyte.

Among aging-related DEGs, we observed that the proportion of tissue-shared DEGs (genes differentially expressed in at least two tissues) in both PAS and PAA lncRNA genes was higher than that in NPA lncRNA genes (two-sided Fisher’s exact test, *P* < 0.001) (Figure 6C), suggesting that pseudogene-associated lncRNA genes respond to aging more broadly across tissues. Considering the broader tissue expression profile of pseudogene-associated lncRNA genes (Figure S6E, S13), we proceeded to investigate the number of tissues in which the genes exhibited aging-related differential expression in each expression breadth group. Both PAS and PAA lncRNA genes were found to exhibit aging-related differential expression in a greater number of tissues, particularly in the group of genes expressed in nine tissues (Figure 6D), suggesting that pseudogene-associated lncRNA genes are more commonly involved in the aging process. One illustrative example of pseudogene-associated lncRNA genes that exhibit aging-related differential expression in multiple tissues is *JPX*, which demonstrates aging-related differential expression in 20 cell types across 8 tissues (Figure 6E). *JPX* has been demonstrated to be directly related to aging and represents a potential therapeutic target for the treatment of age-related atherosclerosis [49]. Overall, these findings indicate that pseudogene-associated lncRNA genes exhibit a positive and extensive response to aging.

To investigate whether pseudogene-associated lncRNA genes show a close association with aging due to transcriptional regulation, we assessed the enrichment of transcription factor (TF) binding sites in the proximal promoters of each category of lncRNA genes in human. Both PAS and PAA lncRNA genes were enriched for the binding sites of aging-related TFs (Figure S14), including NRF1 [50], E2F6, RB1 [51], PML [52], and SREBF2 [53]. Furthermore, the binding sites of longevity factors (FOXO3, SIRT6) [54,55], rejuvenation factor (SOX5) [56], and aging modulators (MTOR, ZBTB7A, CXXC1, SREBF1) [57,58] were also enriched in the proximal promoters of PAS lncRNA genes. Conversely, NPA lncRNA genes were not enriched for the binding sites of these aging-related TFs. These results suggest that pseudogene-associated lncRNA genes are subject to transcriptional regulation of aging-related TFs, thereby closely associating with aging.

To further explore the putative functions of pseudogene-associated lncRNA genes in aging, we conducted GO enrichment analysis based on the guilt-by-association method for aging-related DEGs in each category of lncRNA genes. The aging-related DEGs in both PAS and PAA lncRNA genes were enriched for aging-related biological processes (Figure 6F), including regulation of inflammatory response, NF-kappaB signaling, TOR signaling, response to oxidative stress, cellular senescence, DNA repair, and telomere maintenance. In contrast, the aging-related DEGs in NPA lncRNA genes were not enriched for these aging-related entries. These findings suggest that pseudogene-associated lncRNA genes regulate aging-related pathways in aging.

To investigate whether the close association of pseudogene-associated lncRNA genes with aging is conserved in mouse, we used RNA-seq data from 8 mouse tissues to characterize the aging-related expression changes of pseudogene-associated lncRNA genes (Methods; Table S6). We found that both PAS and PAA lncRNA genes showed a higher proportion of aging-related DEGs and a broader aging-related differential expression compared to NPA lncRNA genes across mouse tissues (Figure S15). To experimentally validate the aging-related expression changes of pseudogene-associated lncRNA genes, we quantified the expression levels of the top 3 PAS and PAA lncRNA genes differentially expressed in the most mouse tissues by quantitative real-time PCR (RT-qPCR). These pseudogene-associated lncRNA genes showed aging-related differential expression in multiple tissues by both RNA-seq and RT-qPCR analysis (**Figure 7**A, B, C; Figure S16). These results confirm the conserved and close association between pseudogene-associated lncRNA genes and aging. Furthermore, we identified two orthologous PAS lncRNA gene pairs and two orthologous PAA lncRNA gene pairs that exhibited aging-related differential expression in both human and mouse (Figure 7D; Figure S17). These lncRNA genes may play a conserved role in aging. Among them, *PVT1* has been shown to be involved in the regulation of aging-related diseases and skin aging [59,60]. We validated aging-related differential expression of mouse orthologs in these orthologous gene pairs by RT-qPCR (Figure 7E). Together, the characterization of aging-related expression changes across tissues and species suggested that the association between pseudogene-associated lncRNA genes and aging is tight and conserved. Therefore, further rigorous studies of the pseudogene-associated lncRNA genes identified in this study will facilitate the discovery of novel aging markers and therapeutic targets for aging-related diseases in the future.

**Figure 7.**
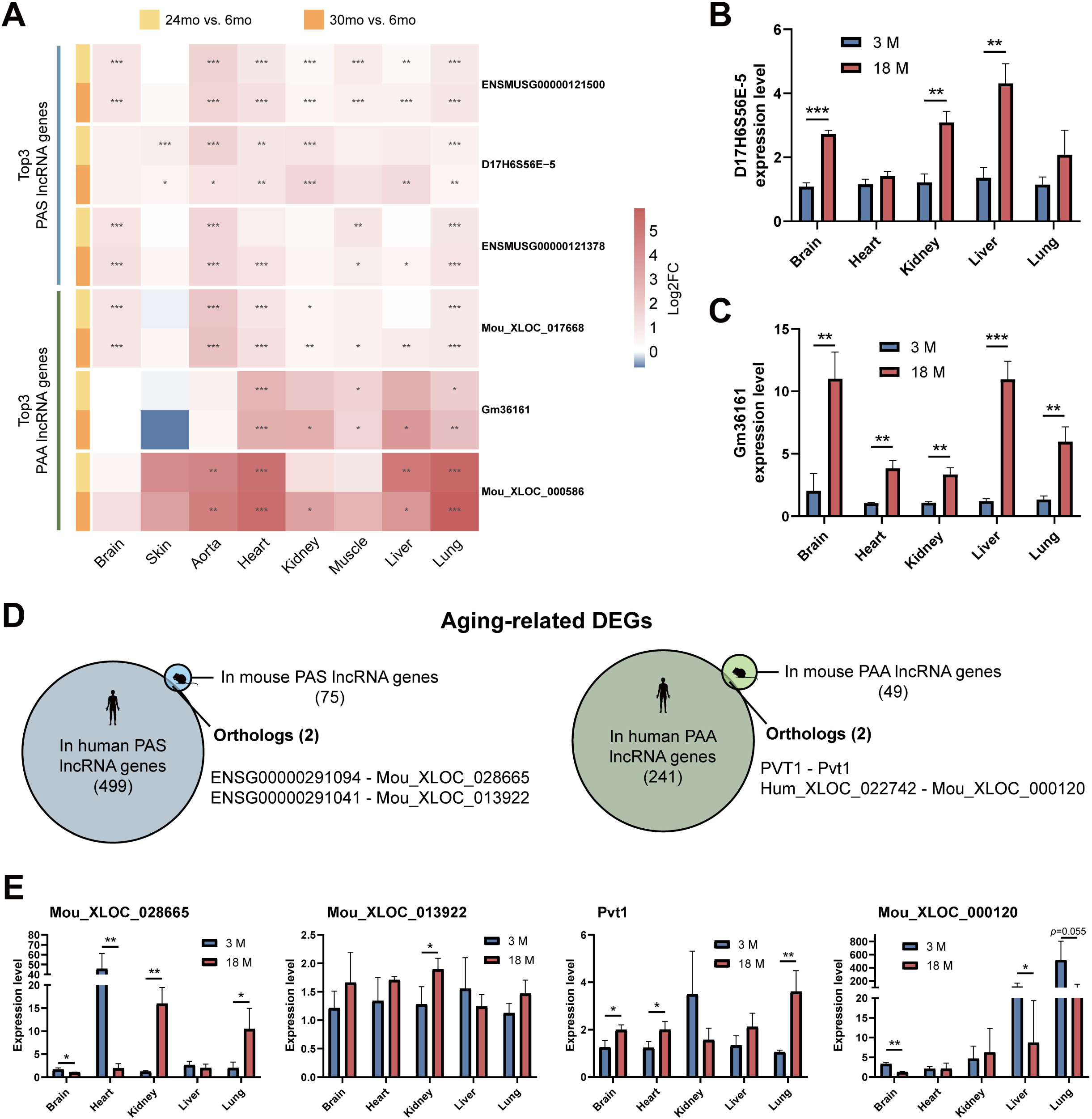
Aging-related expression changes of pseudogene-associated lncRNA genes in mouse. **A**. Aging-related expression changes of the top 3 PAS and PAA lncRNA genes differentially expressed in the most mouse tissues. Statistical significance labels are based on adjusted p-values from DESeq2, \**P* ≤ 0.05, \*\**P* ≤ 0.01, \*\*\**P* ≤ 0.001. **B**. Expression levels of *D17H6S56E-5* in five tissues from 3 month and 18 month old mice, revealed by RT-qPCR. Expression levels are shown as means. Error bars indicate the standard deviation of expression levels. 3 month, n = 3; 18 month, n = 3 mice. \*\**P* < 0.01, \*\*\**P* < 0.001, two-sided Student’s T test. **C**. Expression levels of *Gm36161* in five tissues from 3 month and 18 month old mice, revealed by RT-qPCR. Expression levels are shown as means. Error bars indicate the standard deviation of expression levels. 3 month, n = 3; 18 month, n = 3 mice. \*\**P* < 0.01, \*\*\**P* < 0.001, two-sided Student’s T test. **D**. Orthologous pseudogene-associated lncRNA gene pairs that exhibited aging-related differential expression in both human and mouse. **E**. Expression levels of *Mou_XLOC_028665*, *Mou_XLOC_013922*, *Pvt1*, and *Mou_XLOC_000120* were quantified by RT-qPCR in five tissues from 3 month and 18 month old mice. Expression levels are shown as means. Error bars indicate the standard deviation of expression levels. 3 month, n = 3; 18 month, n = 3 mice. \**P* < 0.05, \*\**P* < 0.01, \*\*\**P* < 0.001, two-sided Student’s T test. RT-qPCR, quantitative real-time PCR.

## Discussion

lncRNA-related research is growing by the day, but there is still an intense debate about the proportion of functional lncRNA genes in the total lncRNA gene collection [6]. Studies of eukaryotic evolutionary dynamics and transcription machinery have shown that lncRNA annotation contain a large number of non-functional lncRNA genes [61]. The co-occurrence of non-functional junk and functional lncRNA genes in lncRNA annotation hinders the broader exploration of functional lncRNA genes. Pseudogenes are often considered as non-functional sequences and thus excluded from functional genomics studies. However, increasing evidence suggests that pseudogenes may be favorable substrates for the origin of functional lncRNA genes [16,19,62].

We identified pseudogene-associated lncRNA genes using genome-wide pseudogenes and a comprehensive lncRNA annotation based on the Ensembl lncRNA annotation, as well as novel lncRNA genes from our home-made Iso-seq datasets and published lncRNA sets from previous studies. More than 40% of pseudogene-associated lncRNA genes are novel lncRNA genes in each species, indicating the significance of a comprehensive lncRNA gene repertoire. Our human lncRNA gene annotation is not as extensive as that provided by the LncBook database, which offers a comprehensive and high-quality collection of human lncRNA genes [63] (Figure S18). However, this limitation arises from the inclusion of eight species in our study. While ensuring systematic construction of the lncRNA gene annotation and comparability across different species, the comprehensiveness of our lncRNA annotation is comparable to that found in recent studies [31,41]. Future enhancements in lncRNA gene annotation across various species will facilitate the identification of pseudogene-associated lncRNA genes.

Our analysis suggests that pseudogenes are favorable substrates for origin of functional lncRNA genes in mammals. This notion encompasses two potential mechanisms: (1) the formation of pseudogenes in intergenic regions, resulting in the origin of novel lncRNA genes, and (2) the integration of pseudogenes into existing genes, leading to the origin of novel chimeric genes that transcribe lncRNAs. Collectively, our findings provide new insights into the origin of lncRNA genes and support the view that new genes can arise from DNA-based or RNA-based gene duplication [64].

Multidimensional evidence supports the enhanced functionality of pseudogene-associated lncRNA genes, including higher evolutionary conservation, higher exon sequence conservation, longer transcripts, more active expression, more active transcriptional regulation, and greater protein-binding capabilities compared to NPA counterparts. In order to provide a more accurate reflection of the influence of pseudogenes on lncRNA genes, a filtering process was conducted on pseudogene-associated lncRNA genes based on stricter overlap length thresholds between pseudogenes and lncRNA genes. We found that pseudogene-associated lncRNA genes showed further heightened functional potential at stricter overlap length cutoffs (Figure S19–22). Taken human PAS lncRNA genes for example, they exhibit invariant evolutionary age distribution and RBP-binding capabilities as well as higher conservation, transcript length, expression levels, and m6A site density at stricter overlap length thresholds (Figure S19). Additionally, transcriptional activity might enhance the identification criteria of pseudogene-associated lncRNA genes. However, the functional potential of pseudogene-associated lncRNA genes is invariant at stricter expression level cutoffs (Figure S23-26). Altogether, these results ensure the reliability of enhanced functional attributes of pseudogene-associated lncRNA genes.

Both PAS and PAA lncRNA genes show an advantage in serving as regulators of gene expression over NPA lncRNA genes. However, PAS and PAA lncRNA genes exhibit distinct functional advantages. Previous studies [62] have implied that PAS lncRNA genes may inherit the RBP binding sites from parent genes of pseudogenes, and our analysis confirmed this view. Therefore, PAS lncRNA genes have greater protein-binding proficiency compared to PAA lncRNA genes. Conversely, PAA lncRNA genes have an advantage in binding to the mRNAs of parent genes of pseudogenes [65,66].

Furthermore, a transcriptome profiling across nine human tissues from young and old age groups at the single-cell level revealed that pseudogene-associated lncRNA genes exhibited prominent and widespread aging-related expression alterations. These pseudogene-associated lncRNA genes are regulated by aging-related TFs and regulate aging-related pathways. Their close association with aging was further confirmed to be conserved in mouse by RNA-seq data and RT-qPCR. Taken together, pseudogene-associated lncRNA genes will constitute a significant resource for the mining of functional lncRNA genes and provide informative insights into human aging biology in the future.

## Materials and methods

### Identification of Pseudogenes

We downloaded genomic data from Ensemble (release-108) for human GRCh38, macaque Mmul_10, mouse GRCm39, rat mRatBN7.2, rabbit OryCun2.0, opossum ASM229v1, chicken GRCg7b, and zebrafish GRCz11, including genome sequences with the repetitive sequences masked, known protein sequences, and genome annotations. The software PseudoPipe used these genomic data to identify genome-wide pseudogene candidates as well as their classification and corresponding parent genes [22]. To eliminate false positives, we performed quality control on pseudogene candidates (identity ≥ 20%; match length ≥ 10% of the query sequence; match length ≥ 30 amino acids; E-value ≤ 1E-5). High-quality pseudogenes that passed quality control were used for downstream analysis.

### Construction of lncRNA Annotation

We complemented the Ensemble lncRNA annotation with long-read transcriptome sequencing data. We first processed mouse, chicken, and zebrafish Iso-seq data generated in our lab. The mouse data have been published [14] and deposited in the GEO database as GSE176018. GSE176018 contains RNA-seq and Iso-seq data for six tissues (including brain, cerebellum, heart, colon, ovary, and testis) from adult C57BL/6J female and male mice. Chicken and zebrafish data include RNA-seq and Iso-seq data for six tissues (including brain, cerebellum, heart, colon, ovary, and testis) from adult female and male chickens and five tissues (including brain, heart, colon, ovary, and testis) from adult female and male zebrafish. Raw Iso-seq data were processed by CCS (v6.2.0) to generate circular consensus sequences (parameters:-j 64--min-passes 1--min-length 300--min-rq 0.9). Circular consensus sequences were processed by lima (v2.3.99) to remove primer sequences to produce full-length sequences (parameters:-j 64--isoseq). Full-length non-chimeric sequences were generated by Isoseq3 refine (v3.5.0) (parameters:-j 64--require-polya). Full-length transcript sequences were generated by Isoseq3 cluster (v3.5.0). RNA-seq data were used by LoRDEC (v0.9) to correct full-length transcript sequences [67]. Corrected transcript sequences longer than 200 were compared to the genome using Minimap2 (v2.26-r1175) (parameters:-t 64 - ax splice--secondary=no-C5-O6,24-B4-uf) [68]. Redundant isoforms were collapsed using the script collapse_isoforms_by_sam.py from cDNA_Cupcake (v29.0.0; https://github.com/Magdoll/cDNA_Cupcake) (parameters:--dun-merge-5-shorter-c 0.85-i 0.95--cpus 28). 5’ degraded transcripts were removed using get_abundance_post_collapse.py and filter_away_subset.py. The above process completed the transcriptome annotation based on long-read transcriptome sequencing data in mouse, chicken, and zebrafish. We obtained human transcriptome annotations based on long-read transcriptome sequencing data from GTEx V9 release.

Bedtools (v2.30.0) was used to select novel genes that did not overlap with Ensemble genes other than pseudogenes from the above annotations based on long-read transcriptome sequencing. We then removed all genes with evidence of coding potential.

CPC2 (v0.1) [69], CNCI (v2) [70], PLEK (v1.2) [71], and LGC (v1.0) [72] were used to detect protein-coding potential. InterProScan (v5.61-93.0) [73] was used to detect transcripts with the Pfam protein domains. Only genes that successfully passed all five filters for all their transcripts were identified as novel lncRNA genes based on long-read transcriptome sequencing.

Next, we obtained high-quality lncRNA gene datasets annotated with short-read transcriptome sequencing data based on multiple developmental stages in seven tissues for seven species (human, rhesus macaque, mouse, rat, rabbit, opossum, and chicken) from the previous study by Kaessmann et al [1]. We also obtained a high-quality lncRNA gene dataset from the FishGET database for zebrafish, annotated based on short-read transcriptome sequencing data [74]. These lncRNA gene dataset were mapped to the genome used in this study using liftOver, and transcripts with changes in transcript structure after mapping were excluded. lncRNAs that overlapped with Ensemble genes other than pseudogenes and novel lncRNAs annotated based on long-read transcriptome sequencing were also eliminated. The remaining lncRNAs were then used as novel lncRNAs annotated based on short-read transcriptome sequencing. These novel lncRNA annotations were merged with the Ensembl annotations as complements.

### Identification of Pseudogene-associated lncRNA genes

We identified two categories of pseudogene-associated lncRNA genes based on two different overlap patterns. Among the total lncRNA genes, we considered genes whose exons overlap with pseudogenes on the same strand as PAS lncRNA genes, and genes whose exons overlap with pseudogenes on the opposite strand as PAA lncRNA genes. All remaining lncRNA genes were classified as NPA lncRNA genes. The data for the two categories of pseudogene-associated lncRNA genes are provided in Table S1 and Table S2.

### Estimation of Enrichment of lncRNA Genes at Pseudogenes

To evaluate the enrichment of lncRNA genes at pseudogenes, Bedtools shuffle (v2.30.0) was used to randomly generate 1000 sets of shuffled regions with the same number and equal lengths as pseudogenes. Shuffled regions were not to be placed in exons of protein-coding genes. Bedtools intersect was used to screen for overlap between pseudogenes or shuffled regions and exons of lncRNA genes. Fold enrichment is a ratio of an observed overlap compared to the expected overlap based on shuffling. The number of pseudogenes overlapping exons of lncRNA genes was calculated as the observed overlap. The mean of the number of shuffled regions overlapping exons of lncRNA genes in each set of shuffled regions was calculated as the expected overlap. The empirical p-value of enrichment is defined as the number of shuffled region sets that show an equal or greater overlap than the observed overlap. To exclude the potential effects of chromosomal biases, we utilized the Bedtools shuffle parameter “-chrom” to make the shuffled regions exhibit the same chromosomal distribution as the pseudogenes. To exclude the effects of repetitive sequences, we downloaded annotations of repetitive sequences from UCSC Genome Browser and excluded pseudogenes and lncRNA genes overlapping with repetitive sequences in the enrichment analysis. We utilized the Bedtools shuffle parameter “-excl” to ensure that the shuffled regions are not placed in repetitive sequences.

### Identification of Homologous lncRNA Genes and Estimation of lncRNA Gene Age

For human and mouse lncRNA genes, we first used liftOver to project their genomic coordinates to other species based on chain files generated by the UCSC Genome Browser (parameter:-minMatch=0.1). We then applied Bedtools intersect to obtain the intersection of lncRNA genes around 10 kb and identify syntenic lncRNA genes across species. For each pair of syntenic lncRNA genes, we used gffread (v0.12.7) to obtain the transcript sequences. Lastz (v1.04.22) was used to obtain the maximum alignment score. We identified the one-to-one orthologous lncRNA genes based on the alignment scores and the reciprocal-best rule. Finally, we used one-to-one orthologous lncRNA genes to infer the evolutionary age of each lncRNA gene based on the phylogenetic relationships of the species in which each lncRNA gene was transcribed, using the parsimony rule. The ages of lncRNA genes with cross-species syntenic lncRNA genes, but no one-to-one orthologous lncRNA genes, were categorized as “ambiguous”.

### Sequence Conservation and Variation Analysis

Genome-wide nucleotide resolution PhastCons scores were downloaded from the UCSC Genome Browser. Average PhastCons scores for exonic regions were calculated using bigWigAverageOverBed. Human common (minor allele frequency (MAF) ≥ 0.01) single nucleotide polymorphism (SNP) data were downloaded from the dsSNP database. The number of SNPs in exon regions was obtained by Bedtools intersect. We obtained the derived allele frequencies of SNPs from phase 3 of the 1000 Genomes Project. To avoid the bias caused by the high mutation rate in the CpG island region on the analysis, we removed the CpG island region in the analysis of SNP derived allele frequency and density. The CADD scores of the SNPs were obtained from the CADD database (https://cadd.gs.washington.edu/).

### Expression Analysis

We downloaded RNA-seq data from E-MTAB-2836 for a total of 200 samples from 32 different human tissues. C57BL/6J male and female adult mouse multiple tissues (including brain, cerebellum, heart, colon, and gonad) RNA-seq data were obtained from a previous study in our laboratory and stored in GSE176018. Raw FASTQ files were quality controlled using FastQC (v0.11.9) and Trim Galore (v0.6.7), and reads were aligned to the reference genome using STAR (v2.7.10b) [75] (parameters:--runMode alignReads--runThreadN 28--outSAMtype BAM SortedByCoordinate -- readFilesCommand zcat--sjdbOverhang 149--twopassMode Basic -- outBAMsortingThreadN 20 --outSAMmultNmax 1--outSJfilterReads Unique -- outSAMattrIHstart 0 --outSAMstrandField intronMotif). Reads with mapping scores (MAPQ) < 20 were filtered out using SAMtools view (v1.16.1) [76]. Finally, gene expression was quantified using featureCounts (v2.0.3) [77] and gene expression was quantified using transcripts per kilobase million (TPM) to normalize gene expression levels. Genes expressed in any sample (TPM ≥ 1) were used for subsequent analysis.

### Characterization of The Epigenome of Pseudogene-associated lncRNA Genes

We obtained human epigenomic data from the NIH Roadmap Epigenomics Mapping Consortium (http://egg2.wustl.edu/roadmap/). We downloaded DHS data for 53 consolidated epigenomes, H3K27ac peak data for 98 consolidated epigenomes, H3K9ac peak data for 62 consolidated epigenomes, H3K4me2 peak data for 24 consolidated epigenomes, H3K4me3 peak data for 127 consolidated epigenomes, fractional methylation data from WGBS for 37 consolidated epigenomes, and chromHMM chromatin status for 127 consolidated epigenomes. In addition, we obtained human housekeeping consensus peaks from a study based on the analysis of 1785 ATAC-seq and 231 scATAC-seq datasets [78]. We calculated the distribution of housekeeping consensus peaks from ATAC-seq, DHS (DNase I hypersensitivity sites), and four histone modification peaks in a total of 4kb regions upstream and downstream of the gene TSSs. The distribution of each peak relative to the transcription start site (TSS) was generated by deepTools [79], and the average fractional methylation of the region around the TSS was calculated by bigWigAverageOverBed. Genome-wide binding sites for TFs based on ChIP-seq data were downloaded from the GTRD database [80]. We identified the proximal promoter of each gene (±500 bp from the TSS) according to the criteria of the FANTOM5 consortium. Among the proximal promoters with active promoter states, we counted the ratio of active promoter chromHMM state regions.

### Characterization of The Features Associated with RNA-protein Binding

N6-methyladenosine (m6A) site data based on m6A-seq and MeRIP-seq data were downloaded from the REPIC database [81]. Transcriptome-wide binding site data for a large number of RNA-binding proteins based on CLIP-seq were downloaded from the POSTAR3 database [82]. We determined the binding ratio (binding length/total exon length) of each RBP on each gene, and RBP richness was defined as the number of types of different RBPs. The Shannon diversity index and Bray-Curtis similarity of RBPs were all calculated using the R package vegan (v2.6-2). In order to evaluate RBP similarity between each category of pseudogene-associated lncRNAs and transcripts from parent genes of pseudogenes, two types of gene pairs were utilized as controls: pseudogene-associated lncRNA genes and random protein-coding genes exhibiting a matched expression level with parent genes of pseudogenes, as well as parent genes of pseudogenes and random lncRNA genes exhibiting a matched expression level with pseudogene-associated lncRNA genes.

### Screening of Co-expressed Genes for lncRNA Genes

We calculated Spearman expression correlations and adjusted p-values (Holm’s multiple comparison test) between each lncRNA gene and each protein-coding gene. To reduce false positives, we downloaded RNA-seq data from E-MTAB-6814 for a total of 313 samples from 7 tissues and 23 developmental stages in human and from E-MTAB-6798 for a total of 317 samples from 7 tissues and 14 developmental stages in mouse. These data, along with those previously used for expression analysis, were used for screening of co-expressed genes for lncRNA genes. Given the accessible chromatin context and the disproportionate RNA expression observed in adult testis, testis samples were excluded from this analysis. We identified co-expressed gene pairs as those with correlation coefficients greater than 0.75 and adjusted p-values less than 0.001. We further analyzed the enrichment of co-expressed genes for each category of lncRNA genes using all lncRNA genes as background. The co-expressed genes significantly enriched (two-sided Fisher’s exact test, *P* < 0.05) by each category of lncRNA genes were analyzed for the enrichment of GO entries. GO enrichment analysis using protein-coding genes co-expressed with at least one lncRNA gene as background. Enrichment analysis was performed using clusterProfiler (v4.8.1) [83].

### Pre-processing and Analysis of sc/snRNA-seq Data

We downloaded sc/snRNA-seq data from GSE261983, HRA000395, GSE183852, GSE167186, GSE180298, GSE243977, GSE136831, GSE255690, and GSE182786 for a total of 85 samples from 9 tissues and 2 age groups (young and old) in human (Table S3). Cell Ranger (v7.1.0) was used to process the raw data and generate UMI count matrices.

The filtered UMI count matrices were imported into R using the Read10X function and then analyzed using the R package Seurat (v4.9.9). After merging Seurat objects for samples from the same tissue, cells with > 500 expressed genes and < 50% of mitochondrial reads were retained. The NormalizeData function was used to normalize the data. The identification of variable genes was conducted using the FindVariableFeatures function, resulting in the selection of 5000 genes for subsequent analysis. Principal component analysis (PCA) was performed using the RunPCA function, and the RunHarmony function was used to remove batch effects from the data. The top 20 PCs were used for clustering and U-MAP analysis. The FindAllMarkers function was used to calculate the marker genes for each cluster. Cell types were identified based on the expression levels of canonical marker genes (Table S4). Genes expressed in each cell type (positive cell rate ≥ 0.01) were used for subsequent analysis.

We performed differential expression analysis between young and old groups using the FindMarkers function in Seurat. Oocytes are not subjected to differential expression analysis, as they are essentially absent from old ovaries. Genes with |Log2FC| ≥ 0.25 and adjusted P-values < 0.05 were defined as aging-related DEGs. Aging-related DEGs in each category of lncRNA genes were analyzed for GO enrichment analysis. We used the guilt-by-association method to associate the lncRNA genes with GO entries based on their co-expressed protein-coding genes.

### Aging-related Differential Expression Analysis in Mouse

We downloaded RNA-seq data from GSE225576 for a total of 96 samples from 8 tissues of 6-, 24-, and 30-months old male C57BL/6J mice. These RNA-seq data were processed as before. Differential expression analysis was performed between 6-and 24-or 30-month-old mice using DESeq2 (v.1.36.0) [84].

### Animals and Treatment

The young (3-month-old) and aged (18-month-old) female C57BL/6L mice used in this study were obtained from the Experimental Animal Center of Huazhong Agricultural University. Animal husbandry and management strictly adhered to Good Laboratory Practice standards. The animals were housed in an SPF-grade facility under controlled environmental conditions, with the temperature maintained at 22 ± 1°C and a 12-hour light/dark cycle. All mice were provided ad libitum access to sterilized feed and drinking water. Animal care and experimental protocols were conducted in accordance with the Guidelines for the Care and Use of Laboratory Animals issued by the National Research Council and approved by the Institutional Animal Care and Use Committee of Huazhong Agricultural University.

### Total RNA Extraction and Quantitative Real-time PCR

Total RNA was isolated using TRIzol reagent (Catalog No. R401-01, Vazyme, Nanjing, China) following the manufacturer’s protocol. cDNA synthesis was performed with HiScript II Q RT SuperMix for qPCR (+gDNA wiper) (Catalog No. R223-01, Vazyme, Nanjing, China) according to the manufacturer’s instructions. RT-qPCR was conducted using ChamQ Universal SYBR qPCR Master Mix (Catalog No. Q711-02, Vazyme, Nanjing, China). Relative RNA expression was calculated using the Ct (2^-ΔΔCt^) method. The expression of target genes was normalized to the expression of *GAPDH*. The RT-qPCR primers are listed in Table S7.

### Ethical statement

The animal experiments were approved by the Animal Ethics Committees of Huazhong Agricultural University and of Institute of Hydrobiology, Chinese Academy of Sciences (approval ID: HZAUMO-2018-070, SYXK2015-0084). All the experiments were performed in accordance with the Good Practice Guidelines for the Use of Animal Research, Experiments, and Teaching.

## Data availability

The raw sequence data reported in this paper have been deposited in the Genome Sequence Archive [85] in National Genomics Data Center [86], China National Center for Bioinformation / Beijing Institute of Genomics, Chinese Academy of Sciences (GSA: CRA023933, reviewer link: https://ngdc.cncb.ac.cn/gsa/s/QV6bw2mz; BioProject: PRJCA037421) that are publicly accessible at https://ngdc.cncb.ac.cn/gsa. The genomic annotations containing all pseudogene-associated lncRNA genes and lncRNA gene classification information used in this study have been deposited in the OMIX, China National Center for Bioinformation / Beijing Institute of Genomics, Chinese Academy of Sciences (https://ngdc.cncb.ac.cn/omix: accession no. OMIX009515, reviewer link: https://ngdc.cncb.ac.cn/omix/preview/CNojc0bX).

## Code availability

All scripts used in this manuscript have been deposited in Github (https://github.com/Z-H-Zhang/Pseudogene-associated_lncRNA_gene) and BioCode (https://ngdc.cncb.ac.cn/biocode: accession no. BT007886).

## CRediT author statement

**Ze-Hao Zhang:** Investigation, Data curation, Formal analysis, Validation, Visualization, Writing – original draft, Writing – review & editing. **Bo-Han Li:** Investigation, Validation, Writing – review & editing. **Yin-Wei Wang:** Investigation, Visualization, Writing – review & editing. **Sheng Hu Qian:** Investigation, Visualization. **Lu Chen:** Methodology. **Meng-Wei Shi:** Software, Visualization. **Hao Zuo:** Investigation, Formal analysis, Visualization, Writing – review & editing, Supervision. **Zhen-Xia Chen:** Conceptualization, Resources, Writing – original draft, Writing – review & editing, Supervision, Funding acquisition. All authors have read and approved the final manuscript.

## Acknowledgements

This study was supported by National Key Research and Development Program of China (2023YFF1001000), The science and technology major program of Hubei Province (2021ABA011), the Foundation of Hubei Hongshan Laboratory (2021hszd012, 2022hszd024, 2022hszd028), and HZAU-AGIS Cooperation Fund (SZYJY2021010). We thank Fan Yao, Shuai Zhang and all members of Chen lab for helpful discussions.

## Supplementary material

**Figure S1 Pipeline for identification of pseudogenes, integration of lncRNA annotations, and identification of the pseudogene-associated lncRNA gene**

**Figure S2 Features of lncRNA genes, pseudogenes, and pseudogene-associated lncRNA genes**

**A.** Transcript lengths of Ensemble lncRNA genes, novel lncRNA genes annotated by long-read transcriptome sequencing, and novel lncRNA genes annotated by short-read transcriptome sequencing. **B**. Exonic sequence conservation for Ensemble lncRNA genes, novel lncRNA genes annotated by long-read transcriptome sequencing, and novel lncRNA genes annotated by short-read transcriptome sequencing. **C**. Maximum expression levels of Ensemble lncRNA genes, novel lncRNA genes annotated by long-read transcriptome sequencing, and novel lncRNA genes annotated by short-read transcriptome sequencing. **D**. Proportion of the three categories of pseudogenes in each species. **E**. Desnsity of the three categories of pseudogenes in each species. **F**. Pearson correlation analysis between the number of pseudogene-associated lncRNA genes and the total number of lncRNA genes. **G**. Bar plot showing the proportion of novel lncRNA genes among pseudogene-associated lncRNA genes.

**Figure S3 Enrichment of sense and antisense lncRNA genes at each category of pseudogenes in each species**

**Figure S4 Functionality characterization of pseudogene-associated lncRNA genes in human and mouse**

**A.** Distribution of PAS lncRNA genes, PAA lncRNA genes, and NPA lncRNA genes in the branches of the mouse phylogenetic tree. Tree tips, lncRNA proportions for each age group. \**P* < 0.05, \*\**P* < 0.01, \*\*\**P* < 0.001, two-sided Fisher’s exact test. **B**. Enrichment of PAS lncRNA genes, PAA lncRNA genes, and NPA lncRNA genes at different evolutionary ages in mouse. A total of 2000 randomly selected NPA lncRNAs were utilized as controls. **C**. Exonic sequence conservation (PhastCons score), for CDS, UTR, PAS lncRNA genes, PAA lncRNA genes, NPA lncRNA genes, and random intergenic regions in mouse. **D**. Exonic sequence conservation (PhastCons score), for PAS lncRNA genes, PAA lncRNA genes, and NPA lncRNA genes in each age group in mouse. **E**. Mean derived allele frequency of non-CpG SNPs segregating in African populations (1000 Genomes Project). Error bars, 95% confidence intervals based on 1000 bootstrap resampling replicates. **F**. CADD scores for SNPs in CDS, UTR, PAS lncRNA genes, PAA lncRNA genes, NPA lncRNA genes, and random intergenic regions in human. CADD, Combined Annotation-Dependent Depletion. **G**. Transcript lengths of protein-coding genes, PAS lncRNA genes, PAA lncRNA genes, and NPA lncRNA genes in mouse. **H**. Average number of exons in transcripts of protein-coding genes, PAS lncRNA genes, PAA lncRNA genes, and NPA lncRNA genes in human and mouse. Error bars, 95% confidence intervals based on 1000 bootstrap resampling replicates.

**Figure S5 Maximum expression levels of protein-coding genes, PAS lncRNA genes, PAA lncRNA genes, and NPA lncRNA genes in various tissues in human.**

**Figure S6 Expression profiles of pseudogene-associated lncRNA genes in human and mouse**

**A.** Maximum expression levels of protein-coding genes, PAS lncRNA genes, PAA lncRNA genes, and NPA lncRNA genes in mouse. **B**. Expression proportions of protein-coding genes, PAS lncRNA genes, PAA lncRNA genes, and NPA lncRNA genes at a series of expression thresholds in mouse. **C**. Mean expression proportions of protein-coding genes, PAS lncRNA genes, PAA lncRNA genes, and NPA lncRNA genes in various tissues in mouse. Error bars indicate standard deviation of the proportion of expression. **D**. Maximum expression levels of protein-coding genes, PAS lncRNA genes, PAA lncRNA genes, and NPA lncRNA genes in various tissues in mouse. **E**. Number of tissues with expression of protein-coding genes, PAS lncRNA genes, PAA lncRNA genes, and NPA lncRNA genes in human and mouse.

**Figure S7 RBP-binding capabilities and putative functions for each category of lncRNA genes**

**A.** Shannon diversity index of RBPs bound to protein-coding gene transcripts, PAS lncRNAs, PAA lncRNAs, and NPA lncRNAs in human. **B**. Density of m6A sites in protein-coding gene transcripts, PAS lncRNAs, PAA lncRNAs, and NPA lncRNAs in each expression level in human. **C**. Richness of RBPs bound to protein-coding gene transcripts, PAS lncRNAs, PAA lncRNAs, and NPA lncRNAs in each expression level in human. **D**. Shannon diversity index of RBPs bound to protein-coding gene transcripts, PAS lncRNAs, PAA lncRNAs, and NPA lncRNAs in each expression level in human.

**A. E**. Number of RBPs enriched by PAS lncRNAs and PAA lncRNAs. **F**. Number of co-expressed genes enriched by each category of lncRNA genes at each p-value level in human. A total of 2000 randomly selected NPA lncRNA genes were utilized as controls.

**B. G**. Number of co-expressed genes enriched by each category of lncRNA genes at each p-value level in mouse. A total of 2000 randomly selected NPA lncRNA genes were utilized as controls. **H**. GO enrichment analysis to annotate the function of NPA lncRNA genes in mouse. **I**. GO enrichment analysis to annotate the function of PAS lncRNA genes in mouse. **J**. GO enrichment analysis to annotate the function of PAA lncRNA genes in mouse. **K**. GO enrichment analysis to annotate the function of PAA lncRNA genes in human.

**Figure S8 UMAP plots showing the annotated cell types in each tissue**

Cell types are indicated by the different colors.

**Figure S9 Expression proportions for protein-coding genes, PAS lncRNA genes, PAA lncRNA genes, and NPA lncRNA genes in each cell type of each tissue**

**Figure S10 Expression proportions of protein-coding genes, PAS lncRNA genes, PAA lncRNA genes, and NPA lncRNA genes in each tissue**

**Figure S11 Number of cell types with expression of protein-coding genes, PAS lncRNA genes, PAA lncRNA genes, and NPA lncRNA genes**

**Figure S12 Proportions of up-regulated and down-regulated genes for protein-coding genes, PAS lncRNA genes, PAA lncRNA genes, and NPA lncRNA genes in each cell type of each tissue**

Two-sided Fisher’s exact tests were conducted to compare pseudogene-associated lncRNA genes with NPA lncRNA genes. \**P* < 0.05, \*\**P* < 0.01, \*\*\**P* < 0.001.

**Figure S13 Among aging-related DEGs, proportions of tissue-specific genes (Expressed in only one tissue) for protein-coding genes, PAS lncRNA genes, PAA lncRNA genes, and NPA lncRNA genes**

\**P* < 0.05, \*\**P* < 0.01, \*\*\**P* < 0.001, two-sided Fisher’s exact test.

**Figure S14 Enrichment of TF binding sites for each category of lncRNA genes in human**

The representative TFs are shown. A total of 2000 randomly selected NPA lncRNA genes were utilized as controls. TF, transcription factor.

**Figure S15 Aging-related expression changes of pseudogene-associated lncRNA genes across eight mouse tissues**

**A.** Proportions of aging-related DEGs for protein-coding genes, PAS lncRNA genes, PAA lncRNA genes, and NPA lncRNA genes in each tissue. **B**. Odds ratios and confidence intervals (95%) between the proportions of aging-related DEGs in each category of lncRNA genes in eight tissues. \**P* < 0.05, \*\**P* < 0.01, \*\*\**P* < 0.001, two-sided Fisher’s exact tests were performed for comparisons between each category of lncRNA genes. **C**. Among aging-related DEGs, proportions of tissue-shared aging-related DEGs for protein-coding genes, PAS lncRNA genes, PAA lncRNA genes, and NPA lncRNA genes. **D**. Number of tissues in which the genes exhibited aging-related differential expression for protein-coding genes, PAS lncRNA genes, PAA lncRNA genes, and NPA lncRNA genes in each expression breadth group. The diamond dots represent the average. \**P* < 0.05, \*\**P* < 0.01, \*\*\**P* < 0.001, two-sided Wilcoxon rank sum test.

Figure S16 Expression levels of *ENSMUSG00000121500*, *ENSMUSG00000121378*, *Mou_XLOC_017668*, and *Mou_XLOC_000586* in five tissues from 3 month and 18 month old mice, revealed by RT-qPCR Expression levels are shown as means. Error bars indicate standard deviation of the expression levels. 3 month, n = 3; 18 month, n = 3 mice. \**P* < 0.05, \*\**P* < 0.01, \*\*\**P* < 0.001, two-sided Student’s T test.

Figure S17 Aging-related expression changes for *Mou_XLOC_028665*, *Mou_XLOC_013922*, *Pvt1*, and *Mou_XLOC_000120*, revealed by RNA-seq analysis

Statistical significance labels are based on adjusted p-values from DESeq2, \**P* ≤ 0.05, \*\**P* ≤ 0.01, \*\*\**P* ≤ 0.001.

**Figure S18 Venn plot shows the overlap between LncBook database and our human lncRNA gene annotation**

The pie plot on the right shows the sources of lncRNA genes that are only present in our human lncRNA gene annotation.

**Figure S19 Functionality characterization of PAS lncRNA genes at a series of overlap length thresholds in human**

**A.** Enrichment of evolutionary ages. **B**. Exonic sequence conservation (PhastCons score). **C**. Transcript lengths. **D**. Maximum expression levels. **E**. Density of m6A sites.

**A. F**. RBP richness. **G**. RBP diversity. \**P* < 0.05, \*\**P* < 0.01, \*\*\**P* < 0.001, Wilcoxon rank sum test.

**Figure S20 Functionality characterization of PAA lncRNA genes at a series of overlap length thresholds in human**

**A.** Enrichment of evolutionary ages. **B**. Exonic sequence conservation (PhastCons score). **C**. Transcript lengths. **D**. Maximum expression levels. **E**. Density of m6A sites.

**A. F**. RBP richness. **G**. RBP diversity. \**P* < 0.05, \*\**P* < 0.01, \*\*\**P* < 0.001, Wilcoxon rank sum test.

**Figure S21 Functionality characterization of PAS lncRNA genes at a series of overlap length thresholds in mouse**

**A.** Enrichment of evolutionary ages. **B**. Exonic sequence conservation (PhastCons score). **C**. Transcript lengths. **D**. Maximum expression levels. \**P* < 0.05, \*\**P* < 0.01, \*\*\**P* < 0.001, Wilcoxon rank sum test.

**Figure S22 Functionality characterization of PAA lncRNA genes at a series of overlap length thresholds in mouse**

**A.** Enrichment of evolutionary ages. **B**. Exonic sequence conservation (PhastCons score). **C**. Transcript lengths. **D**. Maximum expression levels. \**P* < 0.05, \*\**P* < 0.01, \*\*\**P* < 0.001, Wilcoxon rank sum test.

**Figure S23 Functionality characterization of PAS lncRNA genes at a series of expression level thresholds in human**

**A.** Enrichment of evolutionary ages. **B**. Exonic sequence conservation (PhastCons score). **C**. Transcript lengths.

**Figure S24 Functionality characterization of PAA lncRNA genes at a series of expression level thresholds in human**

**A.** Enrichment of evolutionary ages. **B**. Exonic sequence conservation (PhastCons score). **C**. Transcript lengths.

**Figure S25 Functionality characterization of PAS lncRNA genes at a series of expression level thresholds in mouse**

**A.** Enrichment of evolutionary ages. **B**. Exonic sequence conservation (PhastCons score). **C**. Transcript lengths.

**Figure S26 Functionality characterization of PAA lncRNA genes at a series of expression level thresholds in mouse**

**A.** Enrichment of evolutionary ages. **B**. Exonic sequence conservation (PhastCons score). **C**. Transcript lengths.

**Table S1 Pseudogene-associated lncRNA genes in eight vertebrates**

**Table S2 Functionality characterization of pseudogene-associated lncRNA genes in human and mouse**

**Table S3 Sample information of the public single-cell/nucleus RNA sequencing data**

**Table S4 Abbreviations, marker genes, and number for each cell type in each tissue**

**Table S5 Aging-related differential expression of pseudogene-associated lncRNA genes in human**

**Table S6 Aging-related differential expression of pseudogene-associated lncRNA genes in mouse**

**Table S7 Primers for quantitative real-time PCR**

